# Live whole-eye, ex vivo imaging and laser-induced micro injury of the corneal basal epithelium and visualization of resident macrophage responses

**DOI:** 10.1101/2021.06.02.446679

**Authors:** Sebastian M.D. Gulka, Ana M. Litke, Kerry. R. Delaney, Robert L. Chow

**Affiliations:** University of Victoria, Department of Biology, Victoria, Canada

## Abstract

**Purpose:** In this paper, we describe a novel live-imaging approach to visualize the short-term response of the mouse cornea to basal epithelial cell damage. Laser scanning confocal microscopy was used to induce precisely-defined, localized corneal basal epithelial cell damage and the live macrophage response to this damage was visualized and analyzed.

**Methods:** Lipophilic fluorescent dyes, SGC5 or FM 4-64, were injected into the anterior chamber of enucleated eyes and imaged live in whole-mount using confocal laser scanning microscopy. Laser-induced damage was performed by focusing onto a defined region of the corneal basal epithelium for a brief period using a high laser power setting and then returning to low laser power for imaging. Eyes from *CX3CR1*^*+/GFP*^ mice were used to observe macrophage responses to laser damage in real-time.

**Results:** SGC5 or FM 4-64 dyes injected into the anterior chamber readily enter the cornea and are taken up by the stromal layer and labeled the outer membranes of corneal epithelial cells and remained stable when visualized using low laser power. Subjecting a defined region of the basal epithelium to high laser power for 1 minute or longer led to a rapid internalization of dye in the exposed basal epithelium cells and overall increase in cellular fluorescence. This change in fluorescence was also accompanied by cell swelling and contraction. Cellular internalization of the non-lipophilic, dye Alexa 647 hydrazide, indicated that membranes were compromised indicating that exposure to high power laser stimulation causes cellular damage to the basal epithelium. Visualization of corneal resident macrophages close to the site of laser-induced damage showed that within minutes, projecting macrophage filopodia extended towards the damaged region at a rate of 0.75µm/min for roughly 40 minutes.

**Conclusion:** We have developed a novel approach to image the live cornea and its response to damage. Laser-scanning confocal microscopy can be utilized to induce localized damage to mouse corneal basal epithelium and elicit a macrophage morphological response. This approach represents a useful tool for studying corneal wound healing and cellular responses to damage using live whole-mount imaging.

## Introduction

The cornea is the transparent, complex arrangement of cells and connective tissue situated at the front of the eye. It serves a dual role in protecting the internal eye from external damage and in providing much of the refracting power of the eye. Because of the cornea’s importance in vision, it is crucial that any damage it sustains is repaired in a manner that preserves its transparency. The corneal wound repair process is mediated by several cell types including keratocytes, epithelial cells, and macrophages (Wilson *et al*., 2001). In some disease states or corneal damage contexts, the wound healing process may cause hypervascularization and other abnormalities resulting in vision loss (Cursiefen *et al*., 2006).

Resident and infiltrating macrophages have been noted to exert several effects following corneal injury. Depending on the stage of the wound-healing process, macrophages can play both an inflammatory and an anti-inflammatory role (Italiani & Boraschi, 2014; Krzyszczyk *et al*., 2018). Loss of either CCR2- or CCR2+ macrophages delays corneal wound healing as detected with fluorescein staining by about 6 hours (Liu *et al*., 2017a). Corneal macrophages play a key role in directly promoting corneal hem- and lymphangiogenesis following injury (Maruyama *et al*., 2005; Hos *et al*., 2016; Kiesewetter *et al*., 2019). Some of the effects of corneal macrophages in wound healing may be indirect. Interleukin-33 released by corneal macrophages is important in the induction of group 2 innate lymphoid cells which function in tissue repair (Liu, *et al*., 2017b). Autonomic nervous system input on macrophages affects their inflammatory phenotype, thus affecting the progression of wound healing (Xue *et al*., 2018).

While our understanding of the role that macrophages play in corneal wound healing is growing, little has been reported on their acute physiologic response to small-scale damage. Previous methods aimed at damaging the cornea have used methods that cause widespread or extensive damage to the cornea including epithelial scraping (Liu *et al*., 2017a), suturing (Kiesewetter *et al*., 2019), and alkali burns (Kim *et al*., 2012). In addition, very few of the published experimental approaches have utilized live imaging to visualize corneal cellular responses to cellular damage.

Here, we report a novel live imaging, whole-mount approach to study corneal wound healing. Our approach utilizes lipophilic fluorescent dye labeling of corneal cell membranes can be visualized using confocal laser scanning microscopy (Litke *et al*., 2018). Focusing within the plane of the basal epithelium using high laser power leads to robust cellular damage to the exposed cells. Using this damage protocol in *CX3CR1*^*+/GFP*^ mice enabled real time visualization of resident corneal macrophages and revealed rapid responses characterized by the formation of pseudopodia-like processes extending towards the site of damage.

## Methods

### Mice

All experiments were performed following approval by the University of Victoria Animal Care Committee in accordance with guidelines set by the Canadian Council for Animal Care. Mice on a 129S1 genetic background mice were used for laser-damage assay testing. *CX3CR1*^*+/GFP*^ mice (Jung *et al*., 2000) were used for live and fixed imaging of *Cx3cr1*-expressing immune cells. All animals were maintained on a 12-hour light/dark cycle and were 3-6 months of age.

### Anterior chamber injection

Mice were euthanized using isoflurane followed by cervical dislocation. Eyes were enucleated and a pilot hole in the outer edge of the cornea was made using a 30-gauge beveled insulin syringe. A 50 µl Hamilton glass syringe (model 705 RN SYR) with a 31-gauge, beveled needle was used to inject substances into the mouse anterior chamber (AC). When drawing up substances, care was taken to not draw up any air bubbles which would obscure imaging. Eyes were imaged within five minutes post injection.

### Confocal Microscopy

For live whole mount imaging, enucleated mouse eyes were placed inside brain buffer (BB; 0.137M sodium chloride, 2.7 mM potassium chloride, 2 mM calcium chloride, 10 mM HEPES buffer, pH 7.4, and 2 mM magnesium chloride)-filled 4 mm diameter wells in a custom-made petri dish (made by pouring a small amount of Sylgard into a 35mm petri dish and embedding the cut ∼5mm end of a 0.6 mL centrifuge tube into the Sylgard before it dried). Eyes were imaged (without a coverslip) using a 60x water dipping lens (NA 1.0, WD 2.0 mm, Nikon NIR APO 60x water objective, Nikon Corp.) For live imaging, z-stacks were set 1.0-1.5 µm apart. A 488 nm laser was used to image GFP and SGC5, 561 nm for FM 4-64, and 640 nm for Alexa 647 hydrazide. Live time-lapse images were imaged at a pixel density of 512×512 with a pixel dwell of 4.8. Imaging was performed using a Nikon C2 confocal microscope (Nikon Corp., Tokyo, Japan).

### Confocal laser-induced damage

2 μL of 1 mM Alexa 647 hydrazide (Cat. number A20502, Thermo Fisher, Waltham, MA) and 4 uL of 1 mM of SGC5 (Cat. number 70057, Biotium Inc., Fremont, CA) was injected into the AC of an enucleated mouse eye. For *CX3CR1*^*+/GFP*^ mice, only 4 μL of 1 mM FM 4-64 (Cat. number T13320, Thermo Fisher, Waltham, MA) was injected. Eyes were then imaged using confocal laser scanning microscopy. The 488 nm or 405 nm laser was used to damage eyes injected with SGC5, and the 561 nm laser of the confocal microscope for eyes injected with FM 4-64. The focus of the laser was set 6 µm above the basal epithelial-stromal interface. The field of view was zoomed in 6x to scan a region of 1600 µm^2^ and the laser power setting was turned to maximum (100.00). For epithelial damage assays, lasers were run for 60 seconds at various laser power settings. For macrophage response experiments, the laser was scanned continuously for 10 minutes at a resolution of 512 x 512 pixels and pixel dwell was set to 4.8 µsec. The area surrounding the damage was observed for 2-2.5 hours after the damage stimulus using a low laser power setting (2.0).

### Image analysis

#### Image Adjustment

Images were analyzed using the Fiji suite of ImageJ (Schindelin *et al*., 2012). This included 3D drift correction for time-lapse images, pixel intensity adjustments, background subtraction, and distance measurements.

#### Quantification of cell dynamics

ImageJ was used to quantify cell dynamics of both steady-state and injury-responding cells. Time-lapse images were first corrected using plugins>registration>correct 3D drift. A maximum projection of the cell(s) of interest was generated. When measuring length change of cell processes, the first timepoint captured was used as the reference position. A perpendicular line was drawn through the base of the process in the initial reference position and a spot was marked in the middle of that line as the origin to measure from. Lengths were measured from the reference point of the specified cell process to the furthest edge of the cell process in the next timepoint.

## Results

### Confocal laser induces basal epithelial cell damage

In order to visualize corneal cell types in live wholemount preparations, we injected the lipophilic dye SGC5 into the anterior chamber of enucleated eyes. (Figure 1A). We found that SGC5 diffused readily through the stroma and labeled the outer membrane of basal epithelial cells (Figure 1A) and corneal wing cells (data not shown). When imaged at low laser power, membrane labeling of the epithelial cells remained stable and unchanged for >30 minutes.

**Figure 1.**
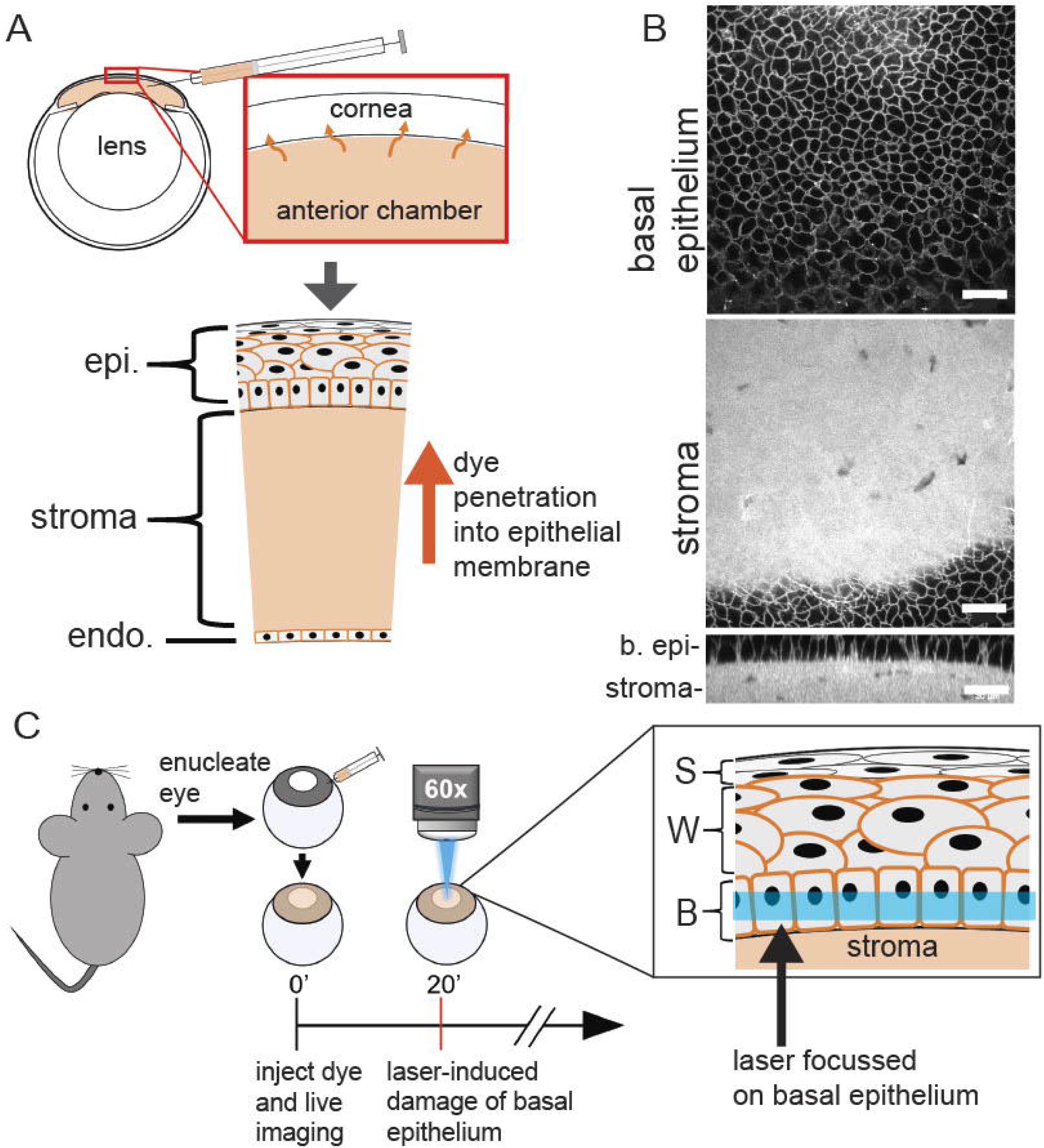
Overview of anterior chamber injection and laser damage protocol. (A) Schematic of cornea and anterior chamber injection. Alexa 647 hydrazide and either SGC5 or FM 4-64 (orange in diagram) were injected into the anterior chamber and diffused into the corneal. SGC5 and FM 4-64 labeled cell membranes while Alexa647 hydrazide was confined to the stroma. (B) SGC5 labeling in the corneal stroma and basal epithelium. (C) Cornea laser-damage protocol. Eyes from euthanized mice were enucleated, injected, and a small region of cornea was imaged at high power for 10 minutes. epi = corneal epithelium, b. epi = basal epithelium, S = superficial epithelium, W = wing cells, B = basal epithelium. Scale bar = 30µm.

To induce damage, corneas were exposed to either the 405 nm or 488 nm laser line set at high power settings and focussed on a single plane set 6 µm above the basal epithelial-stromal interface (Figure 1C). This exposure led to an overall increase in basal cell fluorescence, apparent internalization of fluorescence as early as 3 minutes after the onset of the high laser power (Figure 2). After 10 minutes of exposure, cell swelling followed by contraction was evident in many of the basal epithelial cells early (Figure 2, see cyan colored cell). Similar responses were also observed with as little as 1 minute of exposure to high power laser scanning (Figure 3D). Basal epithelial cells in corneas injected with either FM 4-64 or SGC5 responded identically to laser damage (Figure 3A).

**Figure 2.**
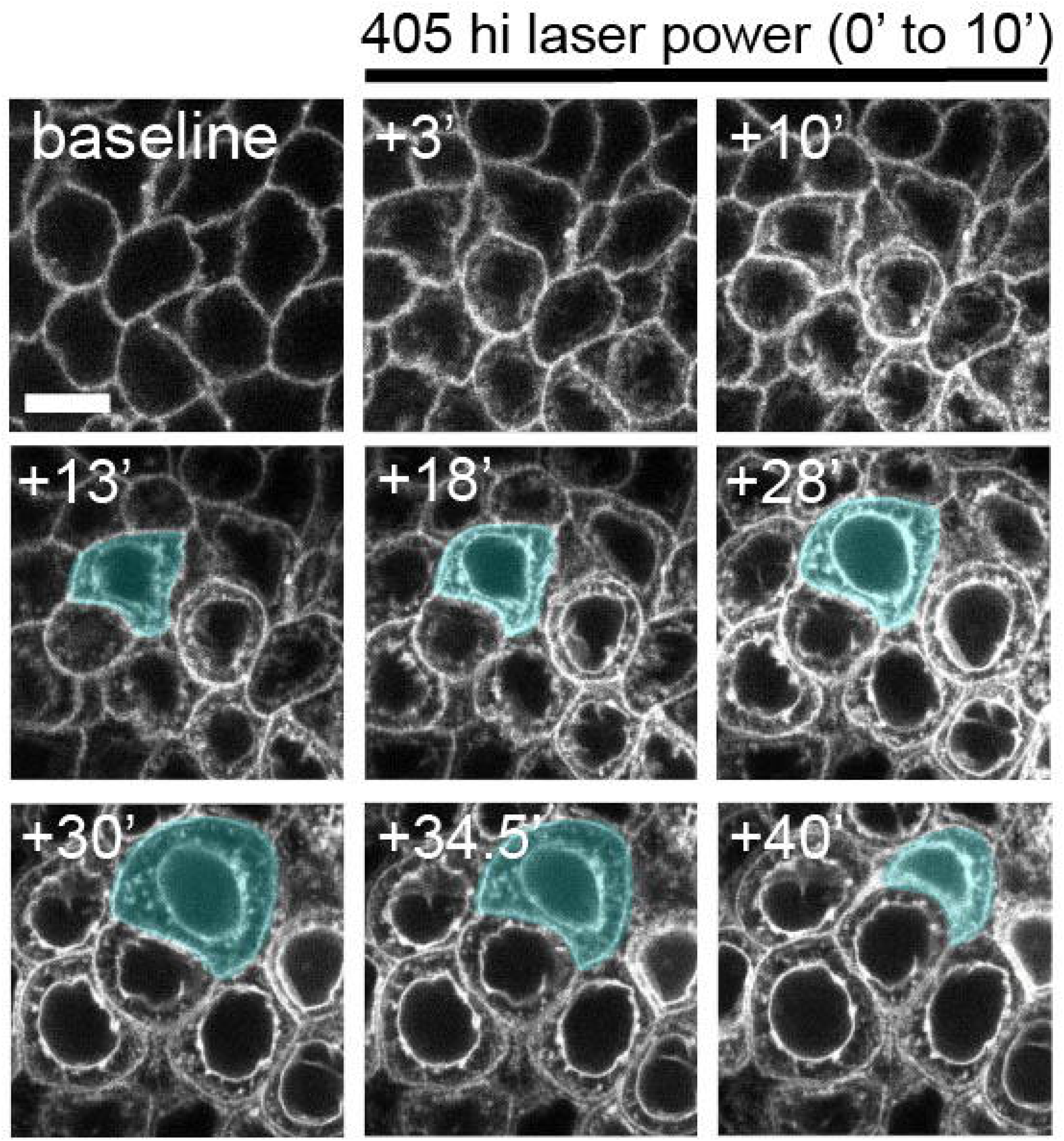
Laser-damaged basal epithelium. Corneas were imaged for 30 minutes with a low-power 488 nm laser which caused no change in membrane dye labeling. The basal epithelium was then co-stimulated with a high-power 405nm laser for 10 minutes. Within 3 minutes, fluorescence was observed within cells, and membrane fluorescence increased. Laser-damaged epithelial cells began swelling followed by contraction within 40 minutes of the onset of the 405nm laser. A cell undergoing swelling and contraction is highlighted In blue. Times are minutes since onset of 405 nm laser. Scale bar = 7.5µm

**Figure 3.**
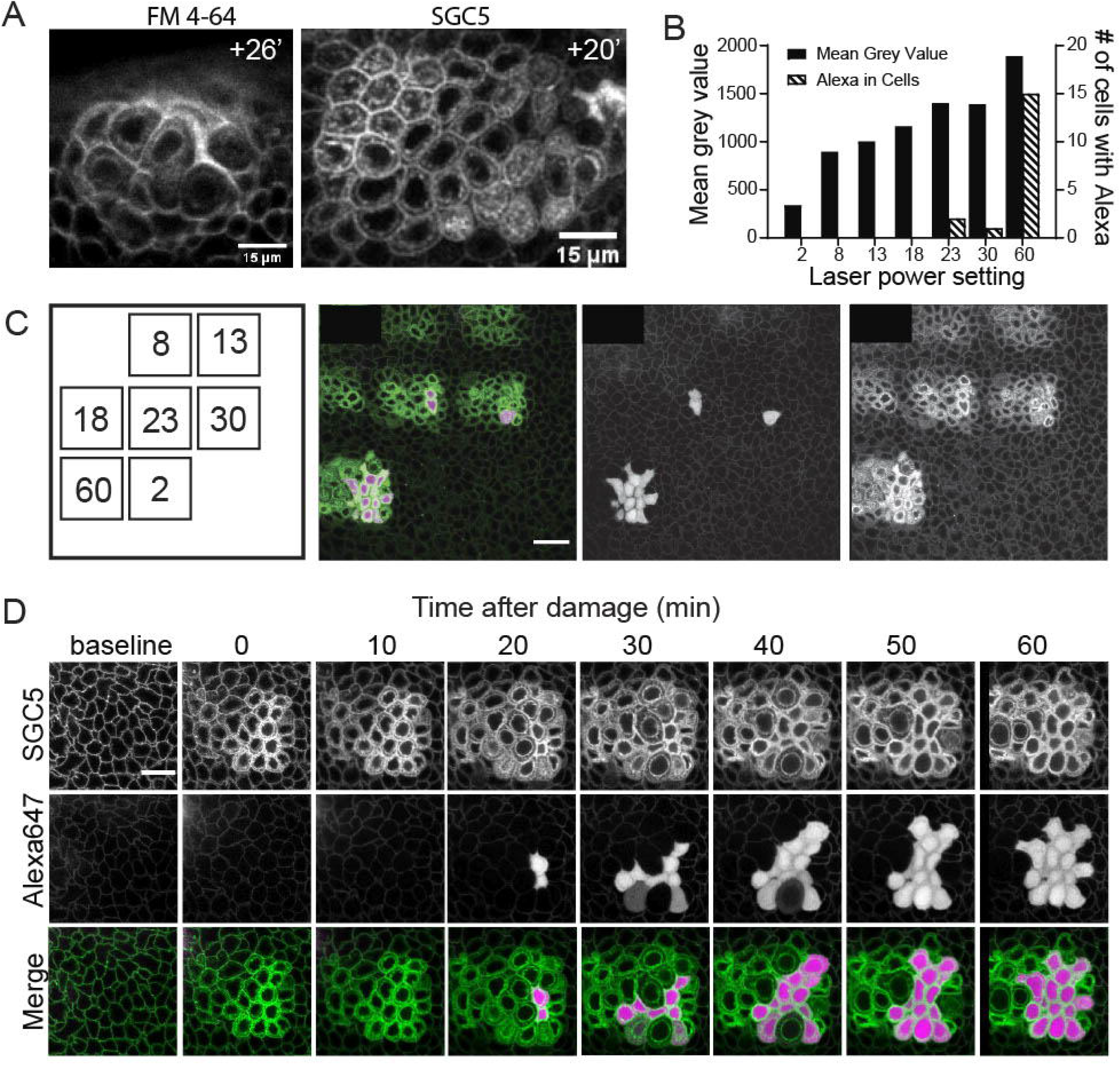
Confocal laser damage effect on basal epithelium of eyes injected SGC5 and Alexa 647 hydrazide dyes. (A) Basal epithelium of corneas injected with FM 4-64 or SGC5 lipophilic membrane dyes behaved similarly after laser damage (B) Alexa 647 hydrazide enters laser-damaged cells at 23.0 laser power setting, while SGC5 intensity increased linearly with laser power. (C) Imaging of corneal basal epithelium from B. The laser power setting for each damaged region is indicated on the corresponding square in the leftmost image. (D) Time-course of corneal basal epithelium damaged with 60 laser-power setting for 60 seconds. Alexa 647 hydrazide penetrates damaged cells 20 minutes-post laser. Scale bar = 30µm (A), 15µm (C).

As the changes in FM 4-64 or SGC5 fluorescence and cell morphology observed following high power laser exposure suggested that the basal epithelium was being damaged, we next wanted to determine whether there were also changes in cell permeability, which is a sign of cell damage (Ünal-Çevik *et al*., 2004; reviewed in Elmore, 2007). To examine this, eyes were co-injected with the hydrophilic dye, Alexa 647 hydrazide. Similar to SGC5, Alexa 647 hydrazide efficiently entered the stroma, but did not penetrate into the epithelium (data not shown). Robust filling of individual basal epithelial cells with Alexa 647 hydrazide was observed in laser-damaged cells after about 20 minutes after the onset of high power laser exposure, suggesting a potential rupture of the cell membrane (Figure 3B,C). A laser-power titration revealed that the intensity of SGC5 fluorescence and presence of Alexa 647 hydrazide in basal epithelial cells increased as the laser power setting increased, with Alexa 647 hydrazide first penetrating into cells at a setting of 23.0 out of 100.0 (Figure 3B). These data reveal a non-lytic membrane damaging of basal epithelial cells by confocal laser, with no obvious damage to the overlying superficial epithelium.

### CX3CR1+ cells response to laser-induced damage

To observe whether the laser-induced basal epithelial damage was sufficient to induce a response in the stromal-localized, resident macrophage population, the enucleated eyes of *CX3CR1*^*+/GFP*^ mice were subjected to the same laser-damage technique mentioned above. Time-lapse videos of undamaged *ex vivo* corneas revealed *CX3CR1*^*+/GFP*^ cells extending and retracting filopodia in a stochastic manner (data not shown). Following laser-induced damage to basal epithelium, nearby macrophages extended pseudopodia to the damaged region (Figure 4). While macrophage pseudopodia exhibited stochastic movements in the naive cornea, following laser damage to nearby basal epithelium, the pseudopodia bud closest to the damage grew markedly in length, while others remained constant, shortened slightly or retract completely to leave the cell in a highly polarized state (Figure 5A,B). There was no evidence that macrophages were migrating toward the site of damage (data not shown). Pseudopodia-like projections from responding macrophages approached and entered the damaged region, reaching full extension roughly 40 minutes after the laser-damage ended (Figure 5C,D). Once pseudopodia reached full extension, morphological changes were not observed in the responding cells for the remaining observation time. Comparing multiple macrophage responses led to a similar trend of an initial filopodia growing phase from 0 minutes to 40 minutes, with an average extension rate of 0.75µm/minute for the first 30 minutes (Figure 5D).

**Figure 4.**
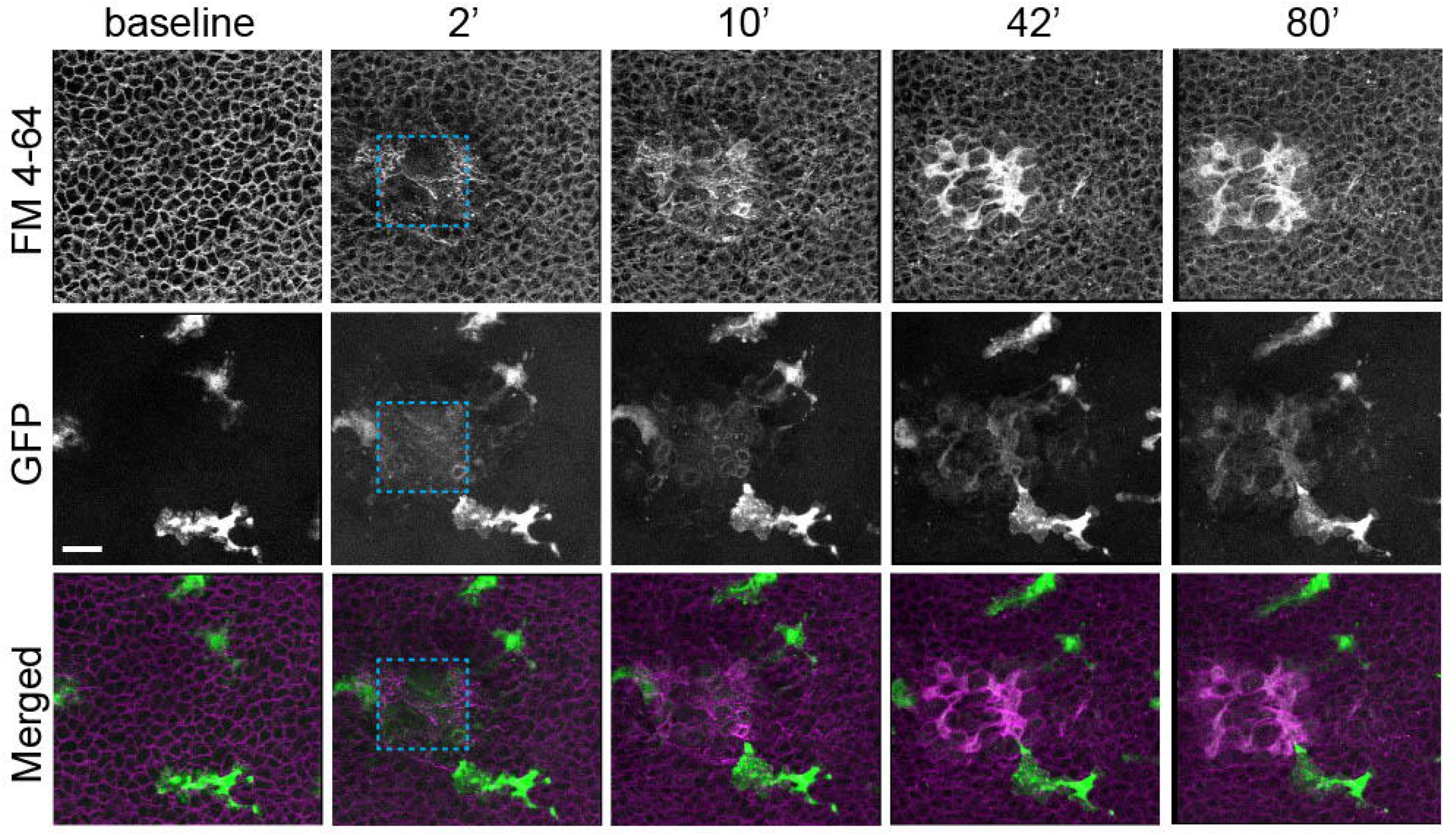
Macrophage response to laser-damaged basal epithelium. (A) When nearby basal epithelium is damaged, macrophages project filopodia to the laser-damaged region (blue square). FM 4-64 (from anterior chamber injection) labels cell membranes, while GFP is from *Cx3cr1:GFP+* cells. Times are minutes after offset of high powered laser exposure. Scale bar = 30µm (A)

**Figure 5.**
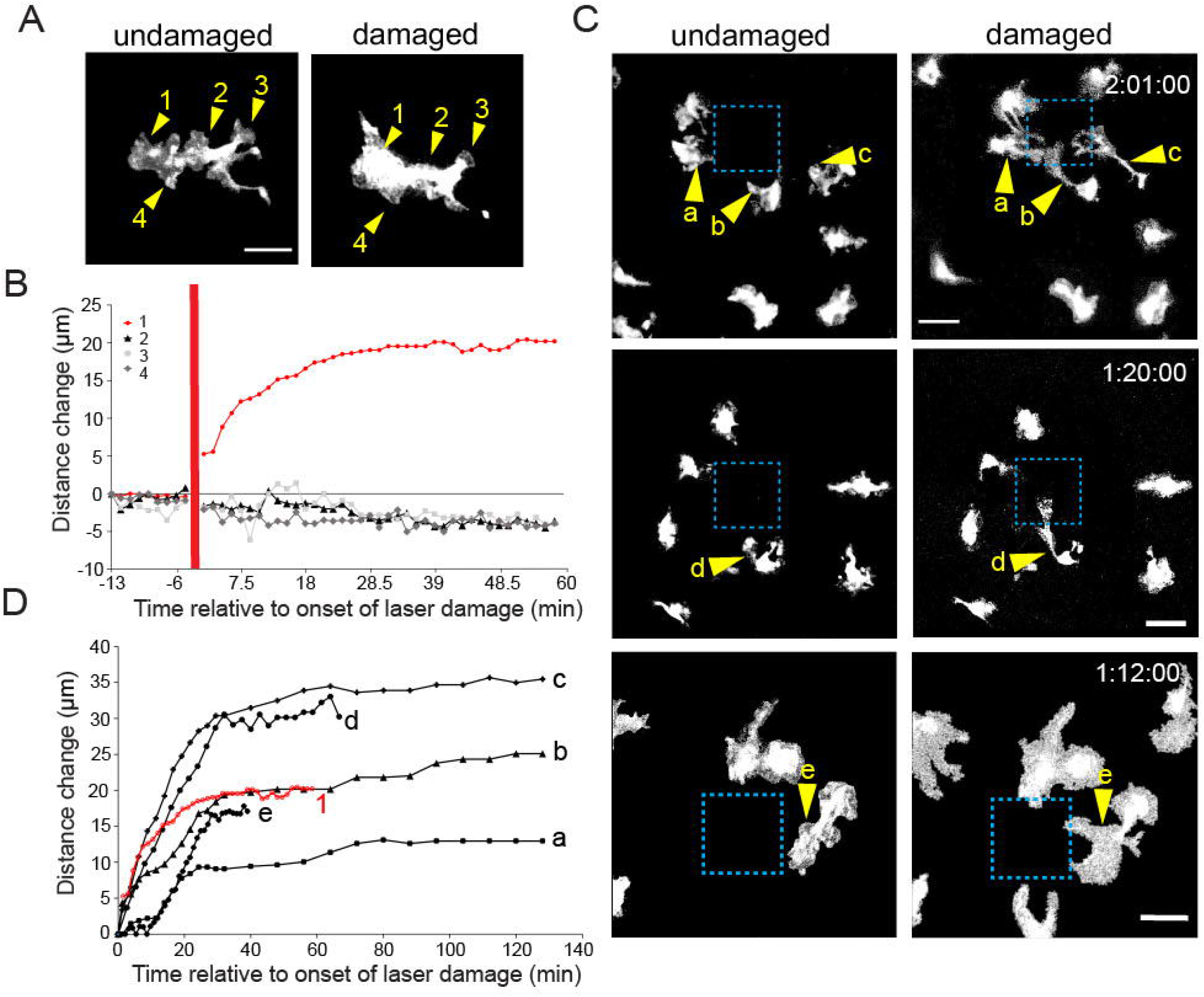
Response of multiple macrophages to laser-damaged basal epithelium. (A) Damage-responding macrophage before and after damage. Labeled processes 1-4 corresponding to the graph in B. (B) Data from the macrophage shown in A reveals that the filopodia closest to the damaged area (process 1) reached full extension after 30 minutes, while other filopodia decreased. Red line indicates time of injury. (C) Live imaging of multiple macrophage responses to laser damage (blue square). Macrophages extended filopodia (yellow arrowhead) to the damaged region. (D) Graph of six responding macrophages from A and C revealed an average extension rate of 0.75µm/min. Times indicated are minutes after offset of high powered laser exposure. Scale bar = 20µm (A) and 30µm (C).

## Discussion

In this study, a novel corneal basal epithelial damage method laser was used, and the live response to damage was captured. The laser-damaged basal epithelium elicited a macrophage response. While lasers have been used in other tissues to cause damage leading to a macrophage/microglial response (Davalos *et al*., 2005; Taylor *et al*., 2018), there have been no studies that investigate the live corneal macrophage response to damage until now. As the damage in our approach was focussed on the basal epithelium, the superficial epithelium was not damaged, and cells were not lysed. Alexa 647 hydrazide penetration into damaged epithelial cells indicates that cell membranes were compromised. Our damage-inducing laser method appears similar to that used by Septiadi *et al*. (2018), who used a confocal scanning laser to cause pulmonary epithelial cell injury. In their case cell death was determined to be the result of apoptosis, with cells maintaining adhesive properties after irradiation.

Both Davalos *et al*. (2005) and Taylor *et al*. (2018) used laser ablation to induce an injury in the mouse brain and observed a microglial response similar to our macrophage response, with responding cells sending processes to the damaged region within minutes of injury. In our study, corneal *Cx3cr1:GFP*+ cells responded to damage on a slightly slower timescale than microglia in the brain, not reaching full extension until 30-60 minutes post-injury in laser-damaged corneas. As excess inflammation may cause corneal defects due to abnormal healing (Matsuda & Smelser, 1973), this may be an intentional way to mitigate those outcomes. The slower response may also be due to the relatively lower density of macrophages in the cornea or the complexity of the stroma preventing efficient diffusion of damage-associated molecular patterns. The corneal macrophage response observed here also appears similar to that seen by Uderhardt *et al*. (2019), with responding cells projecting one or more pseudopodia-like processes towards a laser-damaged damaged region.

While wound-infiltrating macrophages have been noted to play only a minor role in promoting inflammation (Oshima *et al*., 2006), resident macrophages have a significant role in wound evolution. Uderhardt *et al*. (2019) found that resident tissue macrophages in the peritoneum act to cloak small injuries and prevent the influx of neutrophils, which exacerbate tissue damage and inflammation. Given the similarity of our observed macrophage response to theirs, this may be what we observed. Further studies including neutrophil observation are required to confirm this.

Altogether this study shows that fine damage to the corneal epithelium is enough to elicit a stromal macrophage response. Laser-induced injury leads to a localized damage of corneal basal epithelial cells. While the exact nature of this response has yet to be determined, the initial observation of filopodia extension may indicate a cloaking, probing, or phagocytic action. Further studies in the cornea could focus on comparing different types of damage including and beyond what was tested here to determine the threshold for macrophage activation or to see how other immune cells (infiltrating or resident) respond and observing longer timepoints *in vivo* to determine the result of the macrophage response.

## Acknowledgements

This work was funded in part by funding from the National Sciences and Engineering Research Council of Canada and the Canadian Institute for Health Research to RLC.

